# Understanding woody plant encroachment: a plant functional trait approach

**DOI:** 10.1101/2023.07.11.548581

**Authors:** Inger K. de Jonge, Han Olff, Emilian P. Mayemba, Stijn J. Berger, Michiel P. Veldhuis

## Abstract

The increase in the density of woody plants threatens the integrity of grassy ecosystems. It remains unclear if such encroachment can be explained mostly by direct effects of soil conditions and hydrology on woody plant growth or by indirect effects on fire regime and herbivory imposing tree recruitment limitation. Here, we investigate whether woody plant functional traits provide a mechanistic understanding of the complex relationships between these direct and indirect effects. We first assess the role of rainfall, soil fertility, texture, and geomorphology to explain variation in woody plant encroachment following anthropogenically-induced fire suppression across the Serengeti ecosystem. Second, we explore trait-environment relationships and how these mediate vegetation response to fire suppression. We find that woody plant encroachment is strongest in areas with high soil fertility, high rainfall, intermediate catenae positions, and fine soil textures. These conditions promote woody plant communities associated with small stature, small seed sizes, and high recruit densities (linked to a recruitment-stature trade-off). The positioning of species along this ‘recruitment-stature axis’ was found to be the most important predictor of recruitment limitation. Areas that support such plant communities - e.g. mid-catena position - were most sensitive to woody plant encroachment. These findings demonstrate the importance of trait-environment relationships in predicting the impact of human alterations on local vegetation change. Understanding how environmental factors directly (resources) and indirectly (legacy effects and plant traits) determine woody plant encroachment supports the development of process-based ecosystem structure and function models.

## Introduction

Many of the world’s grasslands and savannas are transitioning from herbaceous to woody plant dominance, raising concerns about its potential impacts on ecosystem functioning and services (Eldridge et al., 2011; Sala and Maestre, 2014), including the reduction in productivity (Asner et al., 2004) and the loss of biodiversity (Ratajczak et al., 2012). This widespread phenomenon, known as ‘woody plant encroachment’ (WPE), is influenced by a complex interplay of natural and anthropogenic factors. Regional changes in rainfall patterns (Holdrege et al., 2021), fire suppression (Joubert, Smit and Hoffman, 2012), increased herbivore pressure (Case and Staver, 2017), and global increases in atmospheric CO2 (Stevens et al., 2017; Venter et al., 2018) have all been linked to WPE. Despite the progress in understanding the drivers of WPE, few generalizations emerged that can be used to predict the variation and extent of woody plant increases at local scales, complicating the development of management interventions.

Fire and herbivory have been identified as key processes with strong control over woody cover in savanna ecosystems above a set threshold in mean annual precipitation (MAP) of 650 mm (Sankaran et al., 2005; Lehmann et al., 2014). Fire and browsing herbivores maintain open savannas by preventing the recruitment of woody plants and/or reducing them to smaller size classes (Van Langevelde et al., 2003; Staver and Bond, 2014). This process indirectly favours grasses and so gives rise to a fire-grass feedback, which maintains grasslands even at high rainfall (Beckage et al., 2009; Oliveras and Malhi, 2016). Grazing herbivores can disturb this positive feedback by reducing fuel loads, thereby decreasing fire intensity or suppressing fire altogether (Scholes and Archer, 1997). Sustained grazing over time may even promote an alternative feedback, where woody plants that have escaped the fire and browse traps increase shade, thereby limiting grass growth and fuel loads, promoting further woody cover (Oliveras and Malhi, 2016). These ‘alternative dynamic regimes’ explain the dynamic character of savannas and the strong declines or increases of woody cover at decadal time scales in response to ecological perturbations (Dublin et al., 1990; Sinclair et al., 2008). Currently, increased livestock grazing in and around remaining natural savanna landscapes prevents biomass accumulation in increasingly large areas. The resulting ‘human-driven’ fire suppression has promoted woody cover, especially along the margins of protected areas (Veldhuis et al., 2019), but with very variable outcomes on local scales.

Soil properties and geomorphology mediate the effects that fire and herbivory have on the balance between woody and herbaceous vegetation (Devine et al., 2017) with both direct and indirect causal pathways (Fig. 1, current views). Direct effects include higher tree growth rates at high rainfall and soil nutrient availability, which encourages canopy closure in the absence of fire (Higgins et al., 2007; Murphy and Bowman, 2012). However, the same conditions may increase the productivity of grasses. Whether trees are favoured may thus depend on factors that affect tree-grass competition, such as rainfall variability (including intensity and intermittency) or soil texture (February et al. 2013, d’Onofrio et al. 2015). Environmental factors can additionally influence encroachment through interactions with legacy effects (Fig. 1). For example, high rainfall combined with strong seasonality can increase fire frequencies, suppressing trees at the sapling stage (Higgins et al., 2007). These so-called ‘Gullivers’ can maintain themselves by photosynthesis between fires, but seedling and sapling densities may decrease under intense disturbance regimes or changed growing conditions. The resulting demographic bottleneck may limit woody plant encroachment because there are simply not enough recruits to transition into older-size classes.

**Fig. 1.**
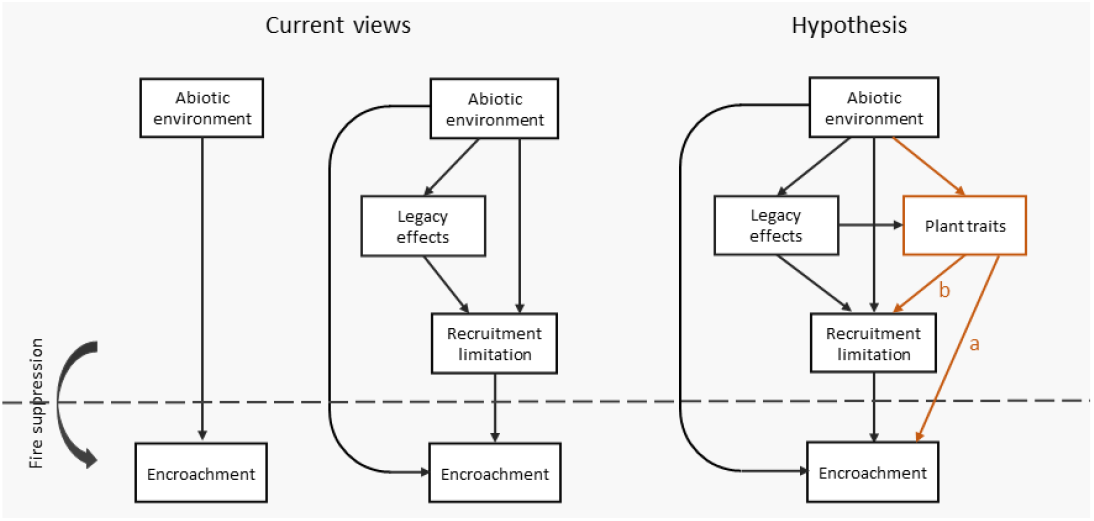
Conceptual models showing current views (left) on how human-driven woody plant encroachment through fire suppression (encroachment) is determined by abiotic environmental factors (abiotic environment), recruitment limitation and legacy effects of disturbance regimes. In this study, we hypothesize (right) that including plant functional traits (plant traits, orange lines) will increase our understanding of the complex relationships between rainfall, soils, fire, herbivory and woody plant encroachment. As outlined in the conceptual models, the indirect effects of the environment on woody encroachment can follow three pathways; through (1) tree recruitment bottlenecks, which may affect encroachment potential, (2) plant traits that control tree growth and recruitment densities and (3) legacy effects of historical disturbance regimes such as fire return interval and the intensity of grazing and browsing (before fire suppression by humans started) which in turn may affect recruitment bottlenecks and plant traits in the community

In this study, we hypothesize that plant functional traits will provide a mechanistic understanding of the variability of WPE (Fig. 1, Hypothesis). Life history traits and functional traits are key concepts in ecology that provide insights into the different strategies that species have evolved to survive and reproduce in varied landscapes and disturbance regimes (Westoby, 1998). Plant traits could therefore help us predict how different plant species will respond to changes in fire regime and herbivore pressure in savannas (Osborne et al., 2019).

Plant size and the leaf economic spectrum (LES) capture the majority of plant trait variation in vascular plants globally (Diaz et al., 2016), providing a useful starting point for examining the relationship between woody plant strategies and WPE. The LES represents a fundamental trade-off between plant species that obtain a slow return on invested carbon (i.e. conservative) and species that gain a fast return (i.e. acquisitive) (Wright et al., 2004). Higher rates of photosynthesis and nutrient uptake enable acquisitive species to quickly pay back the costs of producing new plant material, which should be especially adaptive in productive, high-resource environments. On the other hand, conservative species pay back costs through slow turnover rates of plant material, which generally makes these species more resistant to environmental stressors, but that comes at the cost of reduced growth rates. WPE may be faster in areas that favour acquisitive species (Fig. 1, hypothesis a) because they grow faster and may profit more from the reduction in costs associated with losing perennial structures to fire.

Plant size is strongly linked to demographic metrics because woody plants can only ‘live long’ and attain a large stature if they invest resources into survival (Poorter et al., 2008; Ruger et al., 2018). For example, through growing thick lignified stems -resistance to mechanical damage - and extensive root networks -survival during droughts. High survival rates carry the cost of delayed reproduction because fewer resources are available for producing flowers and seeds. By contrast, short-statured trees generally survive poorly but produce more offspring (Ruger et al., 2018). This should make short-statured species better equipped to recruit or colonize into landscapes where conditions have recently changed than large-statured species that have longer generation times and lower fecundity. At the landscape level, environmental conditions that promote plant communities with poor and delayed recruitment may limit the potential of woody plant encroachment (Fig. 1, hypothesis b).

Investigating whether and how woody plant traits in savannas are related and assessing their importance and specific roles in mediating vegetation response to alterations in fire and grazing regimes is high on the research agenda (Osborne et al., 2019). Although trait-based approaches have emerged as a promising way to understand mechanisms that structure plant communities (Ackerly and Cornwell, 2007; Sonnier et al., 2010), attempts to use these traits to understand local vegetation changes in response to anthropogenic change have so far been limited. Here, we explore the indirect effects of environmental gradients on WPE via their effect on woody plant traits. Specifically, we ask 1) to what extent community means of the main axes of trait variation (economic spectrum and a size dimension) can be predicted from environmental gradients in MAP, soil properties and geomorphology; 2) how the increase of different species after fire suppression is related to their position along these main axes of trait variation; and 3) to what extent the effect of environmental gradients on woody plant encroachment can be explained via indirect effects on plant traits.

## Material and Methods

### Study area and design

We conducted our work within the greater Serengeti-Mara ecosystem, which encompasses approximately 25000 km2 of Northern Tanzania and Southern Kenya and includes the Serengeti National Park (SNP), adjacent land management units and game reserves in Tanzania and the Masai-Mara reserve in Kenya. There is a prominent rainfall gradient from 650 mm MAP in the South-East to 1200 mm MAP in the North-West part of the ecosystem (Mahony et al., 2021). Precipitation is highly variable but generally peaks during a short rain season in December and the long rain season from March-May. Soils transition from volcanic ash-derived plains in the South-East to granite gneisses in the North (Jager, 1982). Vegetation is influenced by both precipitation and soil type and can be broadly classified into the South-Eastern grass plains, West-central Acacia woodlands, and northern broadleaf woodlands (Sinclair, 1995).

Satellite imagery has revealed substantial illegal grazing by livestock inside protected areas, leading to large-scale fire suppression along the margins of protected areas (Veldhuis et al., 2019). In this study, we quantified woody plant encroachment in vegetation plots (n= 96) inside protected areas subject to at least ten years of this form of human-driven fire suppression across the ecosystem (Fig 2a). Woody stem density in fire suppression plots is compared to stem density in control plots (n=91) that experience regular fire. The study was designed to capture important resource gradients, which may influence plant traits of the community and the overall productivity of savanna vegetation. At the largest spatial scale, vegetation plots are distributed over a rainfall gradient from 670 MAP to 1070 MAP and were located to maximize local variation in soil texture, P, N, and exchangeable bases. Soil texture is an appropriate indicator of the hydraulic properties of the soil, with significant consequences for water availability in savannas (Rodríguez-Iturbe and Porporato, 2007). On local scales, the effect of catenae is included (Fig. 2b). Catenae are a major source of landscape heterogeneity in savanna ecosystems, associated with strong soil differentiation and redistribution of rainfall (Khomo et al., 2011).

**Fig. 2.**
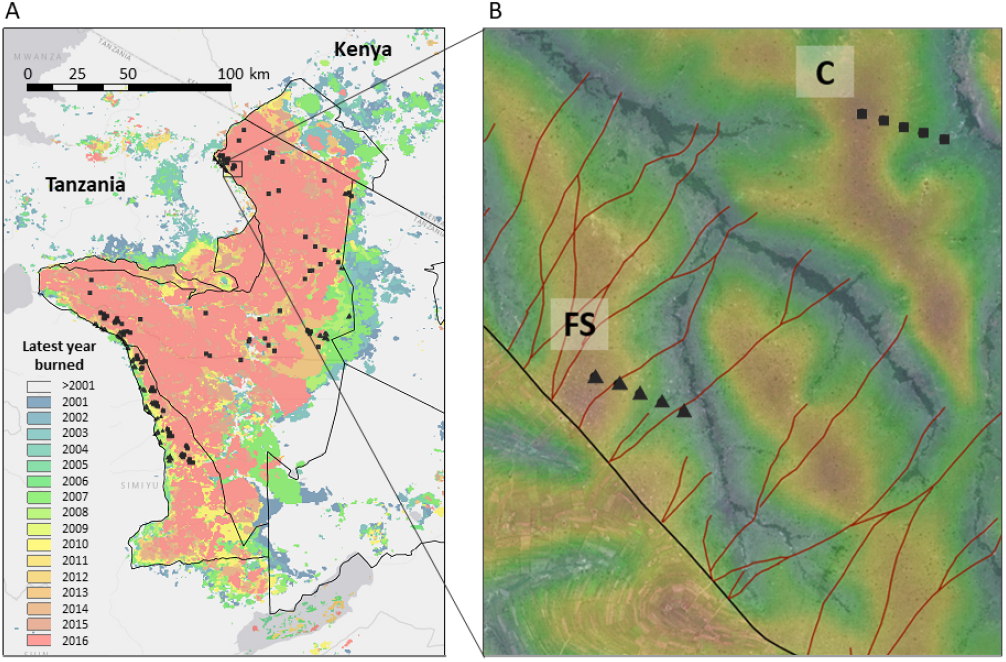
Study design showing vegetation plots (triangles, n=96) along the margins of the ecosystem in zones of human-driven fire suppression and vegetation plots in control zones with no fire suppression (squares, n=91) (A). Zones of high human activity in protected areas are readily visible through ‘livestock’ tracks (red lines in (B). Vegetation plots were kept at least 100 meters from these tracks and 250 meters from the park boundary. The colour scale in (B) reflects the topographic position index (TPI) divided by the standard deviation of the elevation (de Reu et al., 2013) that we used as a measure for the catena position (high to low).

### Woody plant encroachment

Around each central location of a plot, we determined woody stem density within four size classes based on stem diameter at 30 cm height (size class 1: < 1,5 cm stem diameter; size class 2: > 1,5 and < 4cm; size class 3: > 4 and < 15 cm; and size class 4: > 15 cm). Rather than using a fixed-size plot, we sampled the closest individuals to the central location of a plot, resulting in circular plots which are density-dependent in size. This secures a similar sampling effort for different size classes instead of the strong oversampling of saplings and under-sampling of large individuals characteristic of fixed plot sizes. For size classes 1 and 2, we sampled the nearest 20 individuals; for size class 3, the nearest 15 individuals; for size class 4, the nearest ten individuals. Densities were then calculated for each size class by dividing the number of individuals by the (variable) sampled area, assuming a circular plot with the distance to the furthest sampled individual as the radius. As catenae gradients cause relatively fine-scale landscape heterogeneity, we never sampled beyond a radius of 50 meters resulting in treeless vegetation plots in at least some areas. The species identity was determined for each individual. The size-class distribution of woody plants in fire suppression plots versus control plots revealed an increase in saplings (1,5 – 4cm), poles (4 – 10cm), and the smallest mature size classes (up to 20cm) (Fig. S1). Because saplings up to a stem diameter of 4 cm have not yet escaped the fire or browse trap, we used all individuals with a stem diameter between 4 and 20cm in calculating woody plant encroachment.

### Recruitment limitation of tree establishment

Trees can be low in abundance because they hardly recruit (e.g. due to low seed set, seed or seedling predation, or unsuitable environmental conditions for seedlings) or because of low survival from the sapling to tree stage. The first explanation for low tree density, and thus open vegetation in savannas, is called recruitment limitation (Nathan and Muller-Landau, 2000). We used the control plots to assess the effect of environmental factors on seedling densities. The best model, only consisting of abiotic environmental predictors, was used to predict seedling densities in fire suppression plots at the time of fire suppression. Fire suppression plots were classified as being recruitment-limited when the predicted density of seedlings was lower than five individuals per 100 m2. This density would be needed to reach a closed canopy assuming each individual reaches a crown diameter of at least five meters at maturity.

### Disturbance legacies

To explain the variability in WPE across the landscape, we consider the legacy effect of fire and herbivory regimes (before human-induced fire suppression started) on recruitment limitation and plant communities. We used dung counts in transects of 50 m long and 2 m wide to estimate the activity of grazing and browsing wildlife in control plots (see Table S1 for species list and functional guilds). The legacy effect of the fire regime was considered by calculating the proportion of years with fire between 2000 and 2016 using the MODIS MCD64A1 Burned Area Product Collection 6 (Giglio et al., 2009) in control plots.

### Plant traits

For 26 of the most abundant woody species in the ecosystem -accounting for 94 percent of all the individuals observed in our vegetation plots -we measured and collated the following plant traits representing both the carbon-use strategy and life history strategy of plants: leaf nitrogen content (leaf N), leaf carbon to nitrogen ratio (leaf C:N), specific leaf area (SLA), leaf dry matter content (LDMC), leaf thickness (LT), twig dry matter content (TDMC), bark investment, height investment, structural herbivore defence, maximum tree height (Hmax), seed mass, and recruit density. Following the protocol for measuring functional traits by Cornelissen et al. (2003), we collected five mature, sun-exposed leaves of five individuals as much as possible in one region (Seronera, central Serengeti). First, leaf thickness was measured using a micrometer (Mitutoyo) at two standard points for each leaf sample, which were then averaged. SLA was calculated as the ratio of leaf area to leaf dry weight. We weighed the fresh leaves immediately after collection in the field and then oven-dried them at 70°C for 48 hours to obtain the dry weight. LDMC was calculated as the ratio of leaf dry weight to fresh weight. We scanned the leaves using a flatbed scanner to obtain leaf area and analyzed the images using ImageJ software (Abràmoff et al. 2004). The sub-samples were then pooled and ground to fine powder. Carbon and nitrogen contents (percent) were determined by dry combustion using a Flash EA 1112 elemental analyser (Thermo Scientific). Of the same individuals, we additionally collected 2 to 3 stem samples of at least 30 cm from the growing tip to measure TDMC. After collection, the twigs were placed in sealed plastic bags with a damp towel and placed in a cool-box to keep the samples fresh until they could be processed in the lab. To ensure water saturation, the cut end of each twig was placed in water for 6 hours in a dark room, after which the twigs were stripped of leaves and cut to 15cm before measuring their saturated mass. The twigs were placed in a drying oven at 70°C until a constant dry mass was reached and then weighed to determine TDMC.

Bark is an essential component of woody species because it protects and supports the hydraulic tissue (Staver et al., 2020). Woody species in savannas invest in bark to protect meristems from heat and generally have epicormic buds, which allow them to resprout directly from surviving stems (Trollope and Tainton, 1986). Early bark investment may be especially adaptive in low-resource sites because low productivity may restrict the ability of basal resprouting to reoccupy a site after fire (Dantas and Pausas, 2016) and may therefore align with the ‘conservative’ end of the PES. Trees investing in height growth attempt to locate their canopy as quickly as possible above the flame zone before the next fire. They generally rely on high starch reserves in underground crown buds, which enable quick and vigorous post-fire recovery (Clarke et al., 2013), which could relate more to the acquisitive side of the PES. To quantify early bark investment and height investment, we measured the stem diameter, stem height and bark thickness of up to 20 individuals (stem diameter > 1 cm, < 8cm). We measured bark thickness using a standard bark gauge to the nearest millimetre (Haglof, Barktax, Sweden). Thickness and stem diameter were measured between 10 and 50 cm in height, as this is the zone of fire impact on tree stems. The gauge was inserted until resistance by the sapwood was felt. Each species’ early bark investment was quantified by taking the regression slope between stem diameter and bark thickness (Hempson et al., 2014). Diameter-height allometry is informative of how fast a woody plant gains height relative to stem diameter and was therefore used to quantify height investment. We used the same individuals that were measured to quantify early bark investment but discarded individuals which were heavily browsed or damaged. We used additional measurements on undamaged woody plants up to a height of 12 meters. Both height and stem diameter were log-transformed in order to fit linear regressions. For each species, we extracted the regression slope between height and stem diameter as a measure of height investment.

Woody plants in savannas that produce high-quality leaves (acquisitive) are more likely to be structurally defended than species which are low in leaf N (conservative), which are more often chemically defended (Wigley et al., 2018). Structural herbivore defence was classified as being ‘high’ or ‘low’ based on traits that mediate bite size of herbivores: the presence of hooks and leaf-to-spine ratio (Cooper and Owen-Smith, 1986). While cagey architecture is indicative of structural defence as well, it is less suitable in this study as ‘caginess’ is highly variable within species and inducible upon increased browsing levels. Species with either hooks (generally, the Senegalia species) or larger spines than leaves were classified as having ‘high’ structural herbivore defence, while species having neither were classified as having ‘low’ structural defences.

Information on the maximum tree height of different species was extracted from a collection of East African reference sources (Brenan, 1959; Wickens et al., 1973; Sands, 2003; Gillett, 1991), while seed mass was obtained from the Royal Botanical Gardens Kew’s Seed Information Database (https://data.kew.org/sid/) (Liu et al., 2019). Recruit density is often expressed as per capita (i.e., per individual) or per basal area of adult individuals because it standardises the measurement of recruitment across different populations and environments. Measuring standardized recruit density in savanna woody plants is complicated because individuals vary greatly in size and growth rate within species, which makes the determination of established individuals (those that can reproduce) difficult. Moreover, a savanna tree’s size does not necessarily reflect its age, as the tree may have experienced multiple growth cycles interrupted by elephant or fire-induced top kill (Bond and Midgley, 2003). For these reasons, we only consider the recruits themselves. We calculated the density of individuals in size class 1 for each species in the control plots. Because most species in this study occur throughout the ecosystem under different environmental conditions, we consider the maximum recorded recruit density for each species to control for the potential impact of recent fires and/or droughts that could have temporarily reduced recruit density.

### Environmental gradients

We used a modified topographic position index (TPI) algorithm to construct a landform map based on a digital elevation model (DEM) derived from global C-band shuttle radar topographic mission (SRTM), with a 30 m pixel resolution. TPI measures the relative topographic position of a central point as the difference between the elevation at this point and the average elevation within the direct neighbourhood (Weiss, 2001). The Deviation from mean Elevation (DEV), which measures the topographic position of the central point using TPI divided by the standard deviation of the elevation (SD), was found to produce more accurate landform classifications in heterogeneous landscapes (De Reu et al., 2013). As the Serengeti-Mara ecosystem is an example of geomorphological heterogeneity due to volcanism, we performed the landscape classification using DEV with a neighbourhood of 1500 m. The produced map was then used to locate the five plots over a local catenae gradient (Fig. 2b).

We used digital maps from Africa Soils Information Service (AfSIS) to extract soil sand content (w percent), soil clay content (w percent), soil silt content (w percent), soil N (mg kg-1), soil P (mg 100 kg-1), and exchangeable bases (cmol kg-1) for all plots at a resolution of 250 m as measures of respectively soil texture and soil fertility (Hengl et al., 2017). Principal component analysis (PCA) was used to reduce the dimensionality of soil data from sand, clay, and silt and to capture the most variation in the soil texture data. As most water and nutrient uptake occurs in the soil’s top layers (February and Higgins 2010), we used the depth horizon of 5 – 15 cm for both soil properties (sd2). MAP was calculated based on the 0.05º gridded rainfall dataset from the Climate Hazards Group InfraRed Precipitation with Station data (CHIRPS) (Funk et al., 2015).

### Data analysis

We used a phylogenetically corrected Principal Component Analysis (pPCA) to identify patterns and relationships among the 12 plant traits related to plant function and plant size. The PCA approach requires a complete dataset without missing values. In our dataset, 26 out of 312 traits were missing (8,33 percent) (Table S2). We utilized the missMDA package with the ‘regularized’ method to impute missing data. The iterative nature of the algorithm allows for accurate estimation of the missing values, even when the proportion of missing data points is high (> 5 percent). Then, we constructed a phylogenetic tree based on the DNA sequences of the chloroplast gene (cpDNA) maturase K (matK) of the species included in the study using the R package ape. Sequences for each species were obtained from the GenBank® nucleic acid sequence database (Sayers et al., 2020). We used the R package phytools to perform a PCA analysis on the dataset, which accounts for the non-independence of data due to phylogenetic relatedness. The broken stick method was used to determine the number of significant principal components (Legendre and Legendre, 2012).

For each control plot, values along the primary trait axes for each species present were summed and divided by the total amount of species present. To improve the estimate of trait-environment analysis, the dataset was supplemented with 67 extra control plots across the ecosystem for which species composition was measured (but no stem diameter was available), giving a total number of 158 plots. Savanna ecosystems are characterized by a patchy distribution of woody plants with varying densities across the landscape. In such ecosystems, using community-weighted means may result in biased estimates of community trait values, as the presence of a few species (or the absence of rare but functionally important species) with high or low trait values can strongly influence the overall community values. Another form of bias occurs when there is variation in the abundance of different woody plant species within a community and when this variation is not random with respect to the trait values of species (Newbold et al., 2012). Especially trait values associated with a recruitment axis are unlikely to be independent of abundance. Due to these concerns, we used presence-absence data as these are more robust to species abundance and distribution variation and are less sensitive to outliers than CWM.

To assess how woody plant traits, disturbance regimes, woody stem density, and recruitment limitation are related to environmental gradients (MAP, soil texture, soil P, soil N, exchangeable bases, catenae position, and a quadratic term for catenae position), we used a series of Generalized Linear Models (GLM’s) with different error distributions and link functions. We analyzed variation in woody stem densities in control and fire suppression plots across environmental gradients with GLMs assuming a negative binomial distribution with a logarithmic link function. We opted for a negative binomial distribution because of overdispersion and a quadratic relationship between the mean and variance (Ver Hoef and Boveng, 2007). We analysed browser densities in control plots with a negative binomial model. For grazer abundance, we found that the variance is a linear function of the mean, so we used a quasi-Poisson model. Multiple linear regression was used to relate environmental factors and their interactions to the means of the principal component axis, which summarized the variation in multiple plant traits across species. To answer whether plant traits are important in woody plant encroachment, we calculated each species’ percentage increase in stem density in fire suppression plots versus control plots. We related the increase to the species’ position along the main trait spectra. Only species that occurred at least five times in both control and fire suppression plots were included to improve precision. All models were evaluated based on AIC scores, where the least complex but still-sufficient fitting model was selected when, 6.AIC was lower than 2 (Burnham and Anderson, 2004). The best models were used to construct partial residual plots using the Visreg package (Breheny and Burchett., 2017) to visualize the results. Model fit was assessed using D2, the proportion of deviance explained by each model relative to the null model. All models were tested for multicollinearity (variance inflation < 2.0) using the car package (Fox et al., 2012).

Finally, we used structural equation modelling (SEM) to analyze via which pathways environmental conditions influence woody plant encroachment as a result of human-induced fire suppression. SEM aims to understand causal relationships in systems by evaluating the statistical fit between abstract ecological theories and empirical data (Shipley 2000, Grace 2008). We chose the SEM approach because we aim to evaluate both the direct effects of the environment as well as its indirect effects via woody plant traits, legacy effects, and recruitment limitation. We used the conceptual models described in Fig. 1 to develop a series of structural equation models (SEM’s) and tested their fit based on Fishers’ C using the PiecewiseSEM package (Lefcheck, 2016). Because the set-up of this study is based on a space-for-time approach, the conditions (e.g. fire return, recruitment limitation) at the time when fire suppression occurred were predicted based on the best-fitting models for the control plots. For instance, we used the relationship between MAP and fire frequency in control plots to predict the legacy effect of high or low fire in fire suppression plots given the MAP in fire suppression plots. The primary goal of the SEMs was to generate and refine hypotheses about the underlying mechanisms that drive the observed association among abiotic factors, recruitment limitation, traits, and woody plant encroachment. Because the SEM is based on a series of GLMs with different error distributions and quadratic effects of catena, standardized path coefficients are unavailable (Lefcheck, 2016). Instead, we used dominance analysed using the domir package to visualize the relative importance of paths through arrow widths. All analyses were performed in the statistical environment R, version (3.6.1) (R development Core Team 2019).

## Results

### Woody plant encroachment

The density of woody individuals per 100 m2 increased by 287 percent in fire suppression plots (M = 3.89, SD = 5.25) compared to control plots (M = 1.01, SD = 1.12). The density was best predicted by fire suppression, soil N, MAP, soil texture, exchangeable bases, catena position, as well as interaction terms between fire suppression and soil N, MAP, and exchangeable bases (ANODEV, D2 = 0.50) (Fig. 3). Woody plant density increased in response to fire suppression at high soil N (X2 = 10.130, df = 1, P = 0.001), at the higher rainfall end (X2 = 5.815, df = 1, P = 0.016), and higher exchangeable bases (X2 = 4.436, df = 1, P = 0.035) (Fig. 3, significant interactions effects shown by asterisks). Moreover, there are effects of the environment on the density of woody plants, regardless of fire suppression: Soil N (X2 = 4.287, df = 1, P = 0.038) and mid-catena positions (X2 = 4.287, df = 1, P = 0.038) increased density, but particularly at lower rainfall while density was higher at a slightly higher catena position in wetter areas. Finally, density was highest at finer soil texture but mostly in areas with richer soils in terms of exchangeable bases (X2 = 11.363, df = 1, P < 0.001).

**Fig. 3.**
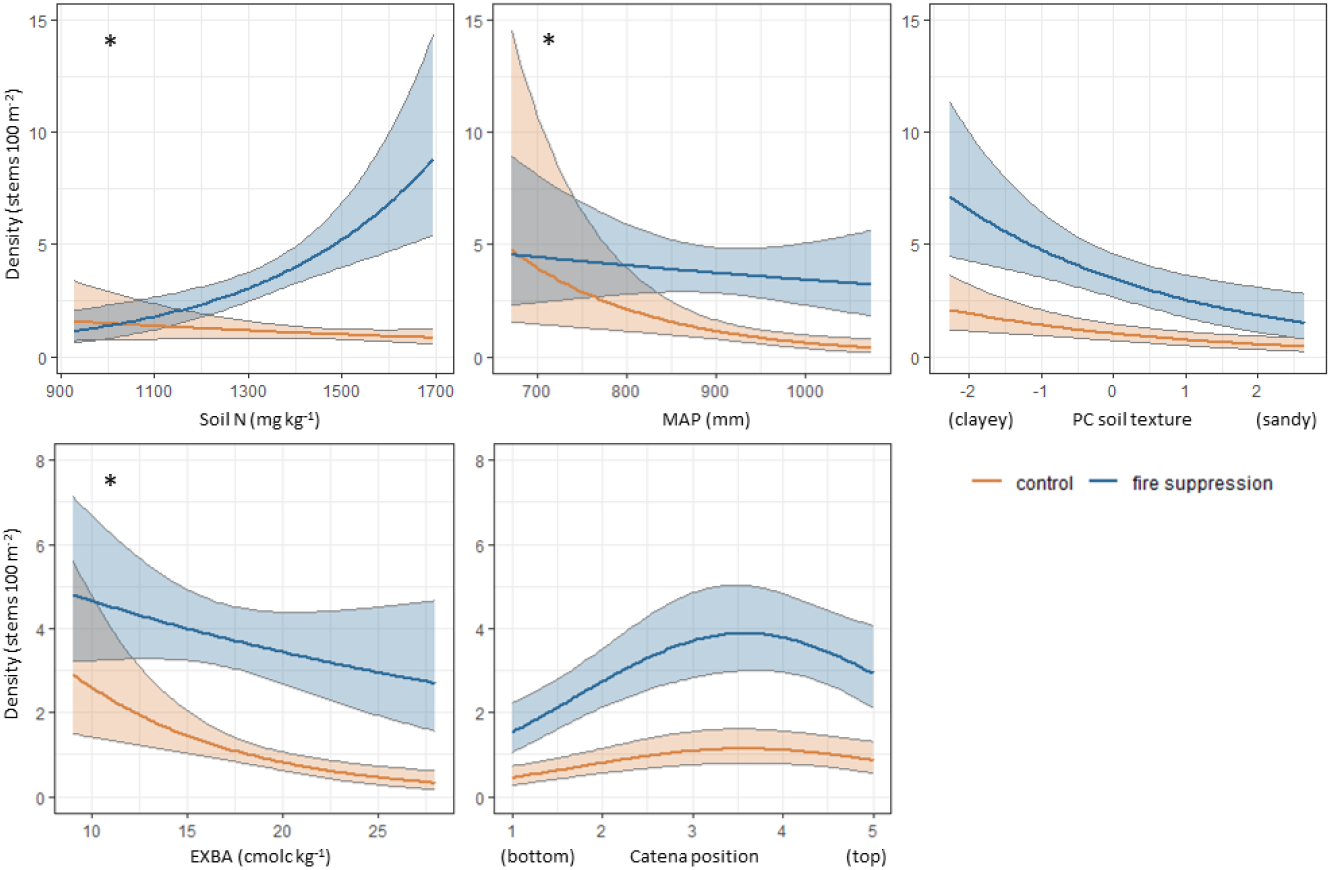
Partial residual plots of the variability in the density of woody plants (size classes poles + young trees, 4 – 20cm) explained by fire suppression, soil N, Mean Annual Precipitation (mm), soil texture, EXchangeable BAses, and catenae position. The partial residual plots are based on the best model and allow for the evaluation of the effect of each predictor on the response variable, given the effect of the other predictors in the model. An asterisk represents significant interactions between fire suppression effects and environmental factors.

### Variation of plant traits

Only the first two principal components (PCs) were retained because they were the only ones that passed the broken stick criterion. Together, they accounted for 67.3 percent of the total variation in the dataset. PC1 explained 46.6 percent of the variation and was primarily associated with plant traits relating to the plant economic spectrum, including leaf traits (Leaf thickness, SLA, LNC, LCN) as well as important fire and herbivore resistance traits (bark investment, height investment, structural defense) (Fig. 4). PC2, accounting for 20.7 percent of the variation, seemed to represent a size dimension (maximum tree height, seed mass) (Fig. 4). While its assignment is somewhat subjective, it appeared that growth-form is associated with the PC2 axis (Fig. 4, see symbols). These results suggest that the first two PCs capture important sources of trait variation in savanna woody plant communities and can be used to investigate the relationships between plant traits, environmental variables, and ecosystem processes. In the following result section, positive PC1 values signify plant communities made up of conservative species, while positive PC2 values signify plant communities with relatively low stature and high reproductive output.

**Fig. 4.**
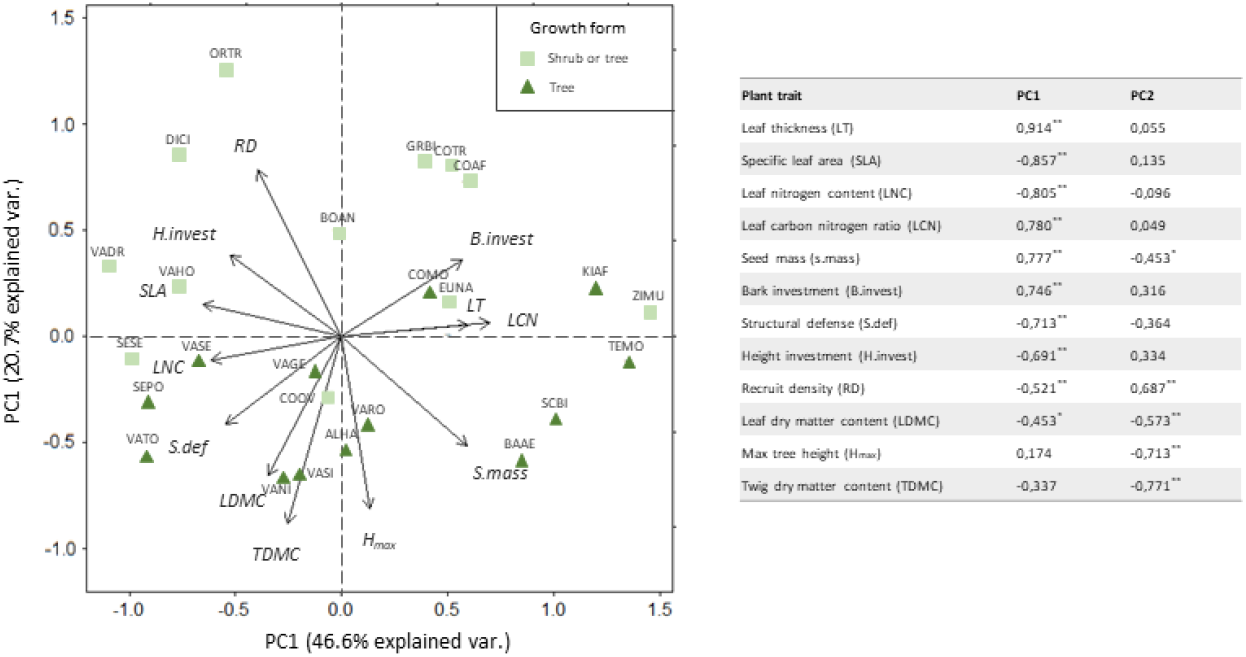
Standardized Phylogenetic Principal Components Analysis (PPCA; first vs second axes) of 26 woody plant species characterized by 12 traits. Plant traits were measured in the Serengeti-Mara ecosystem and derived from the literature. Symbols show species assignment to growth form: The light green squares represent woody plant species that grow as either shrubs or trees, while green triangles represent woody plants that grow as trees (extracted from Brenan (1959), Wickens et al. (1973), Sands (2003), Gillett (1991)). Full species names for the codes can be found in the table. The table gives the bivariate relationships between each plant trait and the scores of PC1 and PC2 across woody plant species. Significant correlations are indicated by boldface and represented with asterisks; *p < 0.05, **p < 0.01

### Trait–environment relationships

Woody plant communities characterized by conservative species (PC1) were positively associated with top catenae positions, high rainfall, and sandy soils (Fig. 5A, F4,153 = 12.78, R2 = 0.25, P < 0.001). Especially at higher rainfall levels, conservative species dominated top catena positions (Fig. 5A, P = 0.048). Communities that have relatively more low-stature species (PC2) were associated with mid-catena positions (quadratic effect), high soil N, low soil P, and high rainfall (Fig. 5A, F5,152 = 14.2, R2 = 0.32, P < 0.001). High-stature species are found relatively more on bottom catena positions (potential drainages) and top catena positions. Dominance analysis revealed that especially the effect of catena (standardized dominance = 0.57) is important in determining the stature of the plant community, followed up by soil N (0.20), soil P (0.13), and rainfall (0.10).

**Fig. 5.**
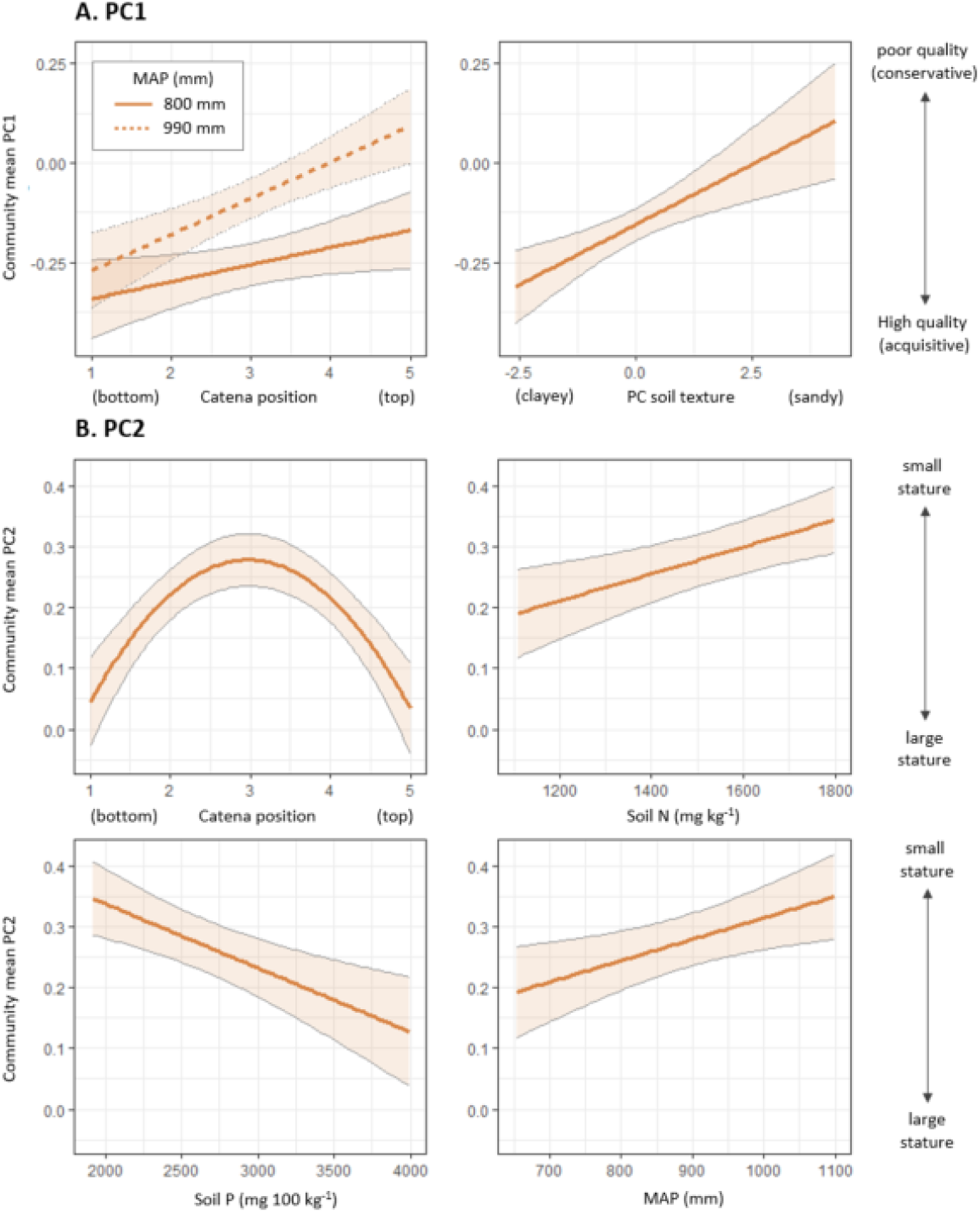
Partial residual plots of the variability in community means of PC1 (A) and PC2 (B) explained by environmental factors. The partial residual plots are based on the best model (according to the lowest AIC) and allow for the evaluation of the effect of each variable on the response variable, given the effect of the other predictors in the model. Note that the catena position of 3 (mid-catena) is used as a condition in constructing the plots showing the effects of soil N, soil P, and MAP on the community mean of PC2 (explaining why the means are relatively high).

### Increase in stem density in relation to traits

The increase in the density of individuals in fire suppression plots compared to control plots was highly variable between species. For instance, the density of Combetrum molle (mostly present in the northern parts of the ecosystem) increased drastically in fire suppression plots compared to control plots (+ 315 percent), suggesting this species acted as an encroacher in response to fire suppression (Fig 6, 7B). In contrast, the density of Balanites aegyptiaca was not different between control and fire suppression zones (Fig. 7B). Species with high values of PC2 (small-stature) increased more in stem density than species with a low PC2 value (high-stature species) in fire suppression plots. A negative exponential relationship fitted the data best (LLR, X2 = 8.834, P = 0.003, Nagelkerke R2 = 0.48). The trait positions along PC1 did not show a relationship with changes in stem density (Fig. 7A).

**Fig. 6.**
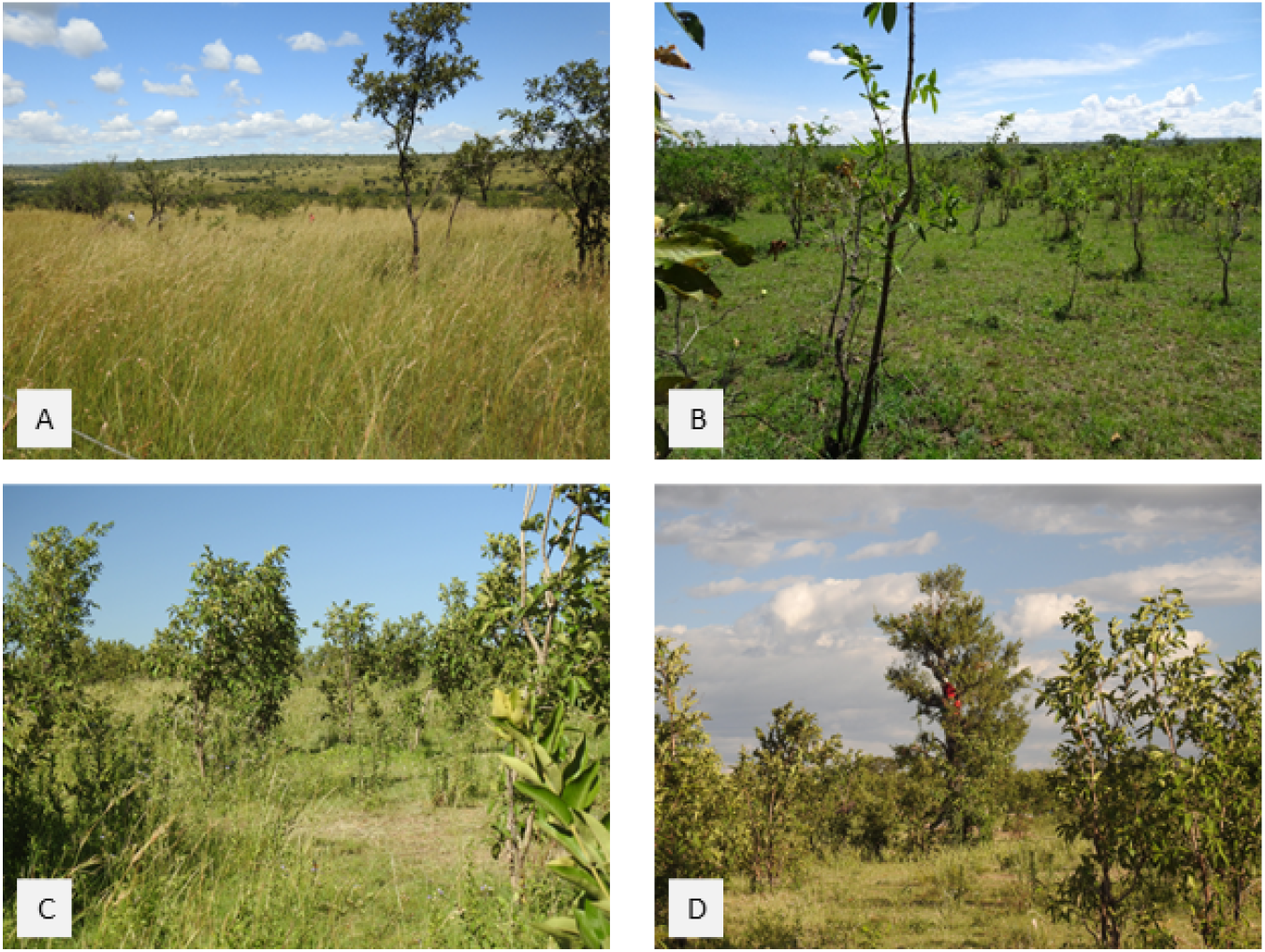
Pictures showing areas dominated by Combretum molle at different stages of fire suppression. Image A shows a ‘control’ plot far from the park boundary with recurrent fire disturbance and high herbaceous biomass. Livestock grazing suppresses fire closer to park boundaries where a dense stand of C. molle saplings arises (B). With continued human-induced fire suppression, these saplings transition to bigger size classes and give rise to woody plant encroachment (C and D). (Photos: Inger K. de Jonge)

**Fig. 7.**
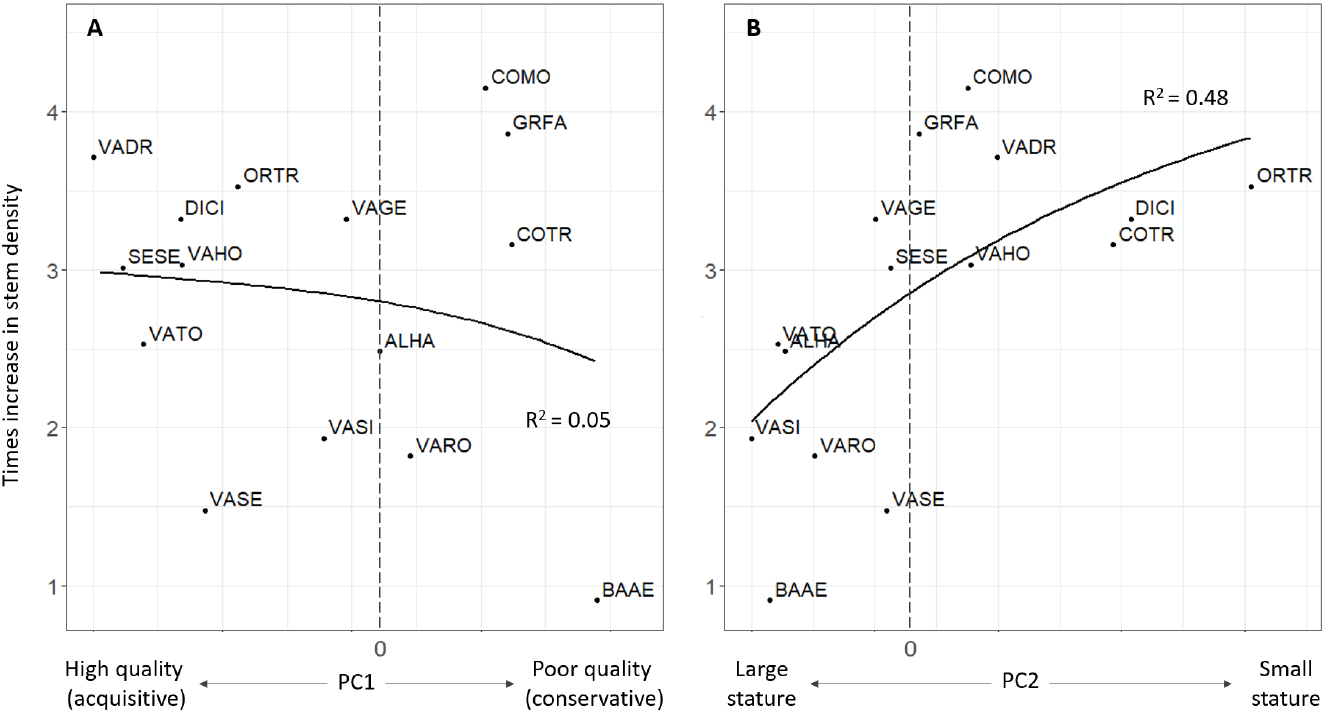
Times increase in the density of woody plant species (size classes poles + young trees, 4 – 20cm stem diameter) in fire suppression plots compared to density in control plots explained by the two main axes of trait variation A. PC1 and B. PC2. Only species that occurred at least in five separate plots in both control and fire suppression conditions were included in the analysis. A value of 1 means no change in density, while four times means a 300 percent increase in stem density. Full species names for the codes can be found in Table S2.

### Alternative pathways for the effect of environmental factors on woody plant encroachment

Three structural equation models were developed and analyzed to compare a model with only direct environmental effects (Fig. 8A), a model also including legacy effects and recruitment limitation (Fig. 8B), and a model including the proposed framework that includes plant traits (Fig. 8C). Because SEM cannot accommodate interaction effects, model A slightly deviated from the results in the first section on woody plant encroachment (Fig. 3). However, soil N, mid catena position, and high rainfall are still linked to higher encroachment (Fig. 8A).

**Fig. 8.**
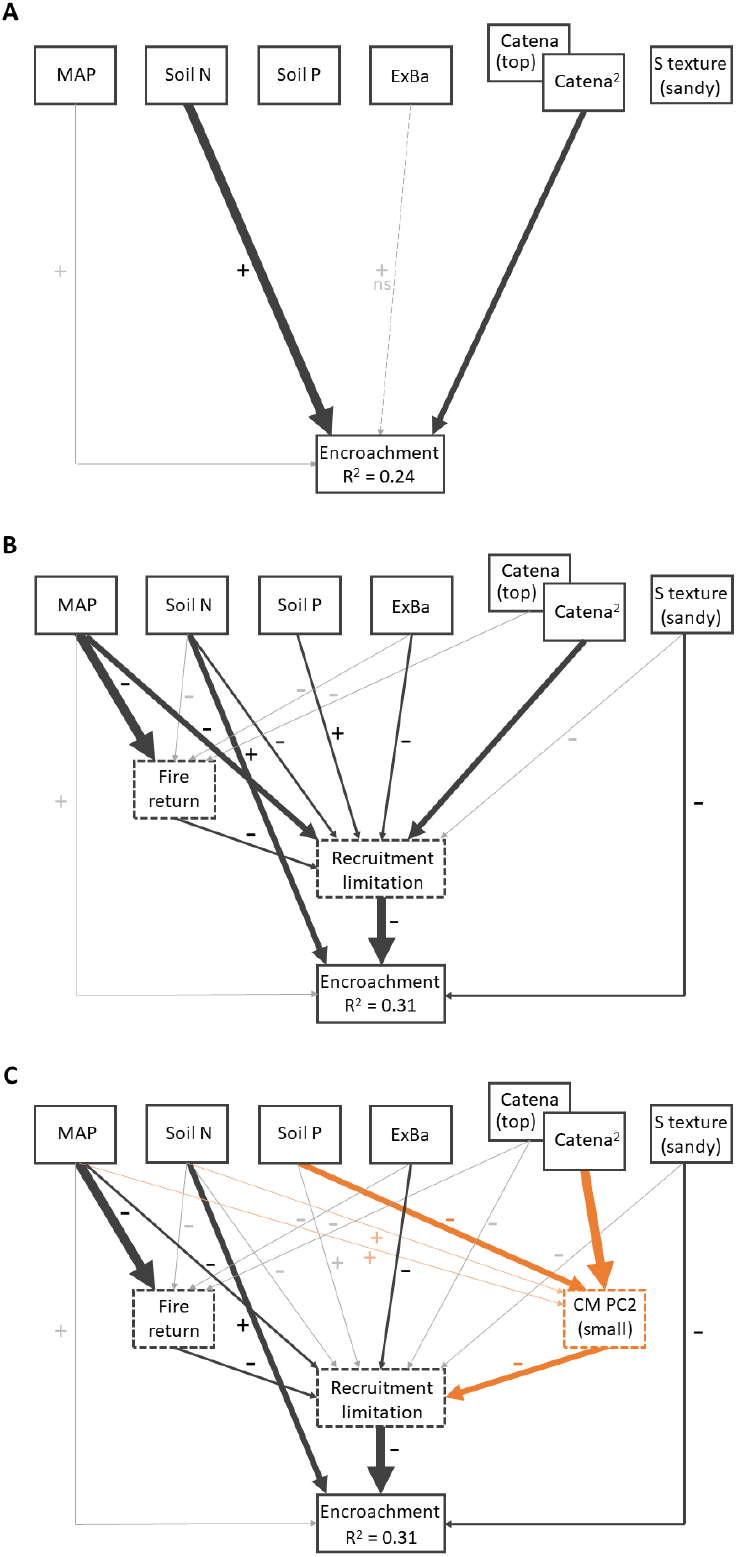
Results of alternative SEM models (A–C) explaining variability in woody plant encroachment following the current views and hypothesis in Figure 1. The top boxes display abiotic environmental factors included in the SEMs; exchangeable bases (ExBa), soil N, soil P, mean annual precipitation (MAP), catena position (reference is top position), and soil texture (reference is sandy condition). Fire return (legacy effect), community mean of PC2 (recruitment-stature axis; reference is ‘small stature’, recruitment limitation, and encroachment are response variables in the SEMs. The sign of the effect is given by -/+, except for the effect of catena2, which is a quadratic effect on encroachment (model A), recruitment limitation (model B), and community PC2 (model C), with the highest levels at intermediate catenae position for encroachment and PC2 and highest levels at the top and bottom positions for recruitment limitation. The orange arrows in C represent the new effects involved with plant traits. R^2^ values associated with the response variables are Nagelkerke’s pseudo-R^2^ and indicate the proportion of explained variation by relationships with other variables. No standardized path coefficients are available as the SEM is based on different Generalized linear models (Lefcheck et al., 2016). Instead, we used dominance analysis of each separate model to visualize the relative importance of relationships through arrow widths.

Seedling densities in control plots increased with soil N and exchangeable bases, especially under lower rainfall conditions (F9,86 = 5.269, R2 = 0.36, P < 0.001). Moreover, seedling density was highest on coarser soil textures but only at bottom to mid-catena positions (P = 0.027). Overall, seedling density was highest at the mid-catena position (P = 0.020). Using this model to predict the density of seedlings in the fire suppression plots, we found that the total number of seedlings was not predictive of encroachment in the SEM analysis, but recruitment limitation (predicted seedlings > 5 individuals/100m2) was (Fig. 8B). This binary classification led to 51 out of 96 fire suppression plots being ‘recruitment limited’ at the time of fire suppression. The mediating effect of recruitment limitation increased the proportion of explained deviance (Nagelkerke R2 = 0.31) of the SEM including legacy effects (Fisher’s C = 7.508, d.f. = 16, P = 0.962) compared to a model with only direct effects of the environment (Nagelkerke R2 = 0.24) (Fig. 8). The SEM additionally included a mediating effect of fire return interval, which predicted that a smaller return interval (i.e. higher fire frequency) leads to recruitment limitation. In this SEM model, it is thus predicted that rainfall has both positive and negative effects on encroachment: directly, high rainfall reduces recruitment limitation, but high rainfall also increases fire frequency, thereby increasing recruitment limitation. Finally, while model A did not include any effects of soil P, model B predicted a negative effect of soil P on encroachment through increasing recruitment limitation (Fig. 8B). Grazing pressure (approximated by dung counts) was highest higher up the catena and at low soil N (ANODEV, D2 = 0.54). Grazing pressure was also highest at low MAP, especially on finer soil textures (P = 0.010). Browsing pressure was highest on soils with fine texture and at low soil P (ANODEV, D2 = 0.46). Moreover, browsing pressure was highest on the top catena at low MAP, while pressure was highest at lower catena positions at high MAP (P = 0.006). Using these models to predict grazing and browsing pressure at the time of fire suppression, we found there was no significant association with either encroachment or recruitment limitation in the fire suppression plots.

The third SEM included the effect of community PC2 (Fig. 5C, Fisher’s C = 12.366, d.f. = 24, P = 0.975). This model predicts that the high encroachment rates at mid-catena positions can be primarily explained by the community and its traits through influencing recruitment limitation (rather than mid-catena directly promoting recruitment). Additionally, model C predicts that the negative effect of soil P on encroachment was mediated by the plant traits of the community through promoting high-stature species. Finally, it predicts that the negative effects of MAP and soil N on recruitment limitation are at least partly mediated by the effects on plant traits (Fig. 8C).

## Discussion

Our study investigated the relationships between woody plant traits in savannas and their importance in mediating woody plant encroachment following fire suppression. We found that woody plant encroachment is most prevalent in areas with higher soil fertility, elevated rainfall, intermediate catenae positions, and finer soil textures (Fig. 3). As expected, at least part of these (abiotic) environmental effects can be explained through indirect effects via recruitment limitation (Fig. 8B, C), an important process controlling woody cover in grassy ecosystems (Midgley et al., 2010). We provide strong support for a causal association between plant traits and recruitment limitation (Fig 1, hypothesis b), where areas that select for small-stature species are less often recruitment limited (Fig. 8C), in turn promoting woody plant encroachment. We will start by discussing the results of the ‘current view’ models (Fig. 1), in which only direct effects of the environment, legacy effects, and recruitment limitations are included. Then, we will discuss how the inclusion of plant traits adds to the interpretation of the data.

### Direct Effects of the Environment on woody plant encroachment

Soil properties played a major role in the increase in the density of woody plants, with the fastest encroachment on clayey, N-rich soils (Fig. 3). Woody plant encroachment - and tree cover in general -is often reported to be higher on coarser soil textures (Sankaran et al., 2008; Case and Staver, 2017), presumably because sandy soils lead to lower fire frequencies and allow for deeper filtration, favouring trees over grasses (Walker and Noy-Meir, 1982). In our study, fire suppression goes hand in hand with increased livestock grazing, which should reduce competition from grasses for water and nutrients. The advantages of fine-textured, N-rich soils (high water-holding capacity, better nutrient supply) thus primarily benefit woody plant growth (Fig 8A). The observed positive effect of rainfall on encroachment can be explained similarly. Without intense competition from grasses, saplings can take full advantage of increased soil moisture, favouring their growth into older size classes. Moreover, rainfall tends to be less variable in higher rainfall areas (Sala et al., 2012), reducing drought-driven mortality of woody plants (Fensham et al., 2009), which could have increased the ability of woody plants to encroach into grassy areas.

### Legacy effects and recruitment limitation

The SEMs that included the mediating effect of recruitment limitation explained more variation than a model that only included direct effects of the environment (Fig. 8B). This is consistent with the idea that woody cover in savannas is a result of episodic but rare recruitment events (Higgins et al., 2000; Bond, 2008), where large numbers of seedlings persist in the understory waiting for a window of opportunity (Balke et al., 2014). Of all the abiotic environmental factors, low rainfall was the most important predictor of recruitment limitation (Fig. 8B). Woody plants have to overcome several demographic hurdles in savannas (Holdo et al., 2014; Midgley and Bond, 2001), of which moisture availability is important in both the seed germination and establishment phase (Loth et al., 2005; Kraaij and Ward, 2006). Wet years combined with frequent rainfall after germination may actually be required for mass recruitment events in savanna ecosystems (Wilson and Witkowski, 1998; Kraaij and Ward, 2006), conditions which may be more often met in wetter areas. High fire frequency was also associated with recruitment limitation (Fig. 8B). Seedlings that have successfully germinated but are still in the early establishment phase (up to 1.5 years) have limited capacity to resprout (Midgley et al., 2010), which should make them vulnerable to annual burns.

Interestingly, the SEMs that fit the data best did not include a mediating effect of grazing or browsing pressure on recruitment limitation. This could have several explanations, the first being that grazers’ adverse and facilitative effects on seedling establishment cancelled each other out. In the earliest recruitment stage, seedlings may be unable to overcome repeated trampling and consumption by bulk grazers (Sinclair, 1995; Hobbs, 2006). However, at a later stage, grazers may facilitate seedlings by reducing grass competition (February et al., 2013; Morrison et al., 2019). Another explanation is that we have only measured dung counts at one moment in time. Herbivore pressure across the landscape can be highly variable in response to stochastic rainfall (e.g. green flushes), and we may not have captured long-term grazing or browsing pressure within our study sites. In a landscape-level experiment, Morrison et al. (2019) found that grass competition was the single most important factor limiting early tree establishment. Although we cannot confirm the causal relationship with increased grass competition in our study, the direct positive effect of soil P on recruitment limitation (Fig. 8B) could represent more intense competition from grasses. Phosphorus tends to be the limiting nutrient for productivity in tropical ecosystems (Vitousek et al., 2010) and because grasses are superior belowground competitors (Cramer et al., 2010), high soil P could lead to competitive exclusion of woody life forms.

### The mediating effect of plant traits

The position of woody plant species along PC2, which linked to maximum height and seed size, was highly predictive of the increase in stem density in response to fire suppression (Fig. 7). The ‘size dimension’ of plant traits has been shown to relate to an important demographic dimension termed the ‘recruitment-stature axis’ in woody plants (Ruger et al., 2018). Large or ‘high-stature’ species maximize canopy performance at the expense of recruitment, while small-statured species maximize recruitment at the expense of survival later on in life (Visser et al., 2016; Ruger et al., 2018). These results thus imply that investment in recruitment rather than survival separates ‘encroaching’ from ‘non-encroaching’ species. Moreover, PC2 was negatively associated with LDMC and TDMC (Fig 4). These traits have been shown to be negatively correlated with potential plant growth rates (Chave et al., 2009), which further explains the ‘encroaching’ habit of species with high values along this recruitment-stature axis (Fig. 5).

The results of the SEMs revealed that the position of species along PC2 is the most important predictor of recruitment limitation (Fig. 8C). The effect of soil P on recruitment limitation is predicted to be primarily indirect through selecting for large-stature species. If soil P indeed intensified grass competition, this indirect pathway raises the question of whether traits associated with PC2 influence the ability of woody species to recruit into dense, high-biomass grasslands. Large-stature species have higher seed mass (Fig. 4), which could give them an advantage in the earliest recruitment phases. Large-seeded species often have improved seedling growth through greater nutrient and sugar reserves, giving them a competitive advantage under stressful conditions (Leishman et al., 2000; Muller-Landau, 2010). This was confirmed in a common-garden experiment by Tomlinson et al. (2019), which showed that seed mass improved the growth of savanna woody seedlings under grass competition. Large-stature species -largely related to the ‘tree’ growth form in savannas -also channel more resources to one big shoot in the recruitment phase (Zizka et al., 2014), which could make them more competitive. However, this competitiveness -increasing the survival of an individual seedling -generally comes at the cost of lower overall reproductive output on the landscape level as survival is traded off against recruitment (Ruger et al., 2018), leading to recruitment limitation in areas that select for large-stature species (Fig. 8C, pathway CM PC2 – recruitment limitation).

Mid-catena positions suffer the most from woody plant encroachment (Fig. 3), which also appears to be mediated by the positive association between mid-catena and small-stature species (Fig. 8C). In a catena sequence, the top and bottom catena suffer least from erosion, have thicker soils, and have a well-developed “A” horizon (Schaetzl and Thompson, 2015). The mid-catena position (sometimes referred to as ‘backslope’) is typically characterized by steep, straight slopes that are often a transitional zone between erosive areas at the ‘shoulder’ of the catena (just below the top) and the bottom of the catena where most deposition occurs (Schaetzl and Thompson, 2015). The moderately developed, shallow soils typical of mid-catena positions may compromise long-term survival (needed in tall-stature species) and promote faster life cycles instead. Indeed, large trees in the Serengeti-Mara are found in well-drained soils (e.g. Vachellia tortilis that use deep taproots to extract water from deep soil layers) or in riparian zones (e.g. Senegalia polyacantha) (Belsky, 1994; Sharam et al., 2009). Low-stature species often have shallower root systems (O’Donnell et al., 2015) and may be better adapted to soil conditions on backslopes. Also, reduced competition from grasses and from large-stature trees may play a significant role: while the top catena has high water infiltration and the bottom catena typically remains wet for prolonged periods, the mid catena may dry out quickly, impairing the productivity of the grass layer. As suggested before by Tomlinson et al. (2019), the grass layer’s biomass dynamics in savannas may thus be largely responsible for the observed trait variation in woody plants, where large-stature species can grow in competition with grasses and small-stature are selected in places where grass production is (temporarily) tempered or is spatially more patchy.

Drier areas were more often recruitment limited, which was also partially mediated by the stature-recruitment axis (Fig. 8C). Seedlings of small-stature species may have a lower tolerance to drought-imposed desiccation through their smaller seed sizes (Daws et al., 2008), limiting their establishment success in areas with infrequent or low rainfall. This could explain why some (otherwise common) small woody species in the Serengeti-Mara, like Ormocarpum trichocarpum and Dichrostachys cinerea, are rare in the driest parts of the ecosystem. The factors that limit the success of small-stature species in drier parts likely extend beyond the impact on the early establishment phase. East-African savannas are highly seasonal, which means that woody plants simultaneously face periodic droughts, fire and herbivory. Low-stature species were associated with low LDMC and TDMC (Fig. 4), likely making these species more vulnerable to physical damage and droughts. D. cinerea, a prominent encroacher of South African savannas as well, was indeed found to be particularly vulnerable to the 2015-2016 drought (Case et al., 2019), and its abundance dropped by 81 percent post-drought (Trotter et al., 2022). Rooting patterns of low versus high-stature species may be important, too: especially shallowly rooting species were found to be sensitive to the 2015-2016 drought (Zhou et al., 2020). Because low-rainfall areas experience droughts more frequently, the number of seed-producing individuals in low-stature species may become the limiting factor rather than establishment limitation. This demonstrates why trait-environment relationships (rather than recruitment limitation alone) are useful for predicting WPE across environmental gradients, especially at local scales.

### Woody plant trait variation in savannas

We used easy measurable plant traits which relate to woody plants’ carbon use strategy and life history strategy and found that the maximum recruit density was positively associated with PC2 (Fig. 4). It is unclear whether these small ‘sizeclass 1’ individuals recruited from seed or whether they were root suckers. O. trichocarpum and D. cinerea are known for their clonal life history and vigorous root suckering (Wakeling and Bond, 2007; Anderson et al., 2015). Root suckering species keep their regenerative organs -including bud banks - and resource reserves underground, making them less sensitive to the impacts of fire (Clarke et al., 2013), explaining why root suckering is quite common in fire-prone savannas, especially in woody plants with unprotected aboveground buds and/or with low bark growth rates (Charles-Dominique et al., 2015; Charles-Dominique et al., 2017). Also, C. molle, the species that increased the most in response to fire suppression (Fig. 6, 7), is known for its root-suckering ability (Bellefontaine, 1997). Establishing whether the dense C. molle sapling stands (Fig. 6) were actual root suckers is difficult as this would have required excavation of the root system. While resprouting through root-suckering is technically not a form of recruitment, we expect that root-suckering is associated with PC2, because it allows small-stature species to persist in the population despite their higher vulnerability to top-kill. A better understanding of which properties associated with the recruitment-stature axis explain the density of trees in encroached areas is needed. Field-based observations point towards specific traits, such as root suckering, but high-quality data on belowground traits has only recently started to emerge (Wigley et al., 2019; Zhou et al., 2020).

## Conclusion

Continued pressure on savanna ecosystems in sub-Saharan Africa requires new approaches with which we can understand the specific effects of human activities in savannas. In this study, we confirm the potential of using a trait-based approach to predict vegetation change in response to anthropogenic impacts (Osborne et al., 2018). Our results show that local variation in woody plant encroachment is strongly associated with the presence of small-stature high-recruitment species, which themselves are likely constrained by competition from grasses, low rainfall, and favoured at mid-catena positions. Because WPE requires management interventions at local scales, understanding the mechanisms that determine the level of encroachment -i.e. plant functional types -will inform management practices (e.g. where to perform preventive burning) and assign resources across large ecosystems (e.g. ranger patrols to reduce livestock grazing).

## ACKNOWLEDGEMENTS

We would like to thank the management and research staff at Serengeti Wildlife Research Center, Tanzanian Wildlife Research Institute, Tanzanian National Park Authority and Tanzanian Commission for Science and Technology for their help and support while undertaking this study. This study was part of the AfricanBioServices project and funded by the EU Horizon 2020 grant number 641918.

