## Supplementary material for "Understanding woody plant encroachment: a plant functional trait approach": Fig. S1; Table S1; Table S2

Appendix S1

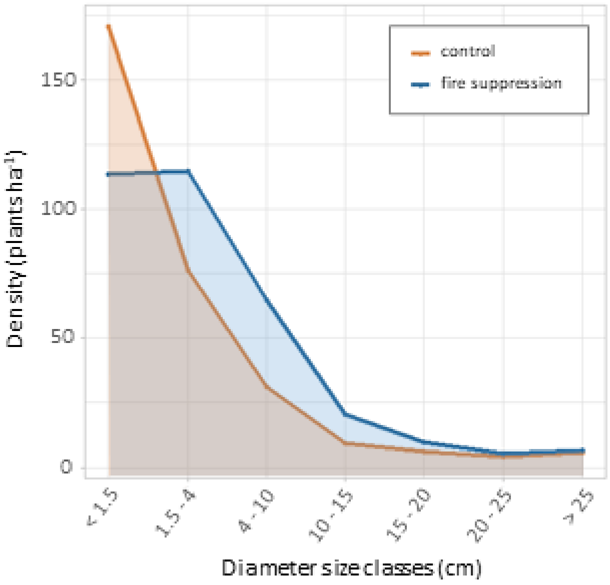


*Fig. S1. Diameter size-class distribution of woody plants in fire suppression plots (blue line and shading) and control plots experiencing regular fire (orange line and shading). Size classes are based on stem diameter at 30cm below ground level.*

*Table S1: List of herbivore species with information on their functional guild (grazer, mixed-feeder, browser)*

| ***Species*** | ***Scientific name*** | ***Functional guild*** |
|---|---|---|
| ***Buffalo*** | *Syncerus caffer* | *Grazer* |
| ***Cattle*** | *Bos taurus* | *Grazer* |
| ***Dik dik*** | *Madoqua kirkii* | *Browser* |
| ***Duiker*** | *Sylvicapra grimmia* | *Browser* |
| ***Eland*** | *Tragelaphus oryx* | *Mixed feeder* |
| ***Elephant*** | *Loxodonta africana* | *Mixed feeder* |
| ***Giraffe*** | *Giraffa camelopardalis* | *Browser* |
| ***Goat*** | *Capra aegagrus hircus* | *Browser* |
| ***Grant's gazelle*** | *Nanger granti* | *Mixed feeder* |
| ***Hartebeest*** | *Alcelaphus buselaphus* | *Grazer* |
| ***Impala*** | *Aepyceros melampus* | *Mixed feeder* |
| ***Oribi*** | *Ourebia ourebi* | *Mixed feeder* |
| ***Reedbuck*** | *Redunca redunca* | *Grazer* |
| ***Roan*** | *Hippotragus equinus* | *Mixed feeder* |
| ***Sheep*** | *Ovis aries* | *Grazer* |
| ***Suni*** | *Neotragus moschatus* | *Browser* |
| ***Thomson's gazelle*** | *Eudorcas thomsonii* | *Mixed feeder* |
| ***Topi*** | *Damaliscus lunatus* | *Grazer* |
| ***Wildebeest*** | *Connochaetes taurinus* | *Grazer* |
| ***Zebra*** | *Equus quagga* | *Grazer* |

Table S2: Table of woody plant species, abbreviations (Code), Leaf nitrogen content (LNC), Leaf C:N ratio (LCN), Leaf dry matter content (LDMC), Leaf thickness (LT), specific leaf area (SLA), Early bark investment (Bark inv), Early height investment (Height inv), Seed mass, Maximum tree height (Max height), Maximum recruit density (Max RD), Twig dry matter content (TDMC), and mechanical defense against herbivores (mech defense) used in the PCA analysis and trait-environment analysis across vegetation plots in the Greater Serengeti-Mara Ecosystem.

| **Species** | **Code** | **LNC (%)** | **LCN** | **LDMC** | **LT (µm)** | **SLA (mm² mg⁻¹)** | **Bark inv** | **Height inv** |
|---|---|---|---|---|---|---|---|---|
| *Albizia harveyi* | ALHA | 2,58 | 18,43 | 0,479 | 208,5 | 10,75 | 0,175 | 0,478 |
| *Balanites aegyptiaca* | BAAE | 2,65 | 16,10 | 0,383 | 349,3 | 6,53 | 0,168 | 0,402 |
| *Boscia angustifolia* | BOAN | 2,39 | 19,24 | 0,397 | 210,8 | 11,10 | | |
| *Commiphora africana* | COAF | 2,00 | 21,45 | 0,359 | 247,1 | 10,08 | 0,249 | 0,476 |
| *Combretum molle* | COMO | 1,65 | 28,12 | 0,441 | 344,6 | 6,37 | 0,164 | 0,570 |
| *Cordia ovalis* | COOV | 2,34 | 19,94 | 0,518 | 190,6 | 8,49 | | |
| *Commiphora trothea* | COTR | 1,69 | 24,42 | 0,346 | 262,8 | 9,02 | 0,222 | 0,599 |
| *Dichrostachys cinerea* | DICI | 2,11 | 21,59 | 0,429 | 144,0 | 14,98 | 0,146 | 0,757 |
| *Euclea natalensis* | EUNA | 2,03 | 22,68 | 0,402 | 272,2 | 8,72 | | |
| *Grewia bicolor* | GRBI | 2,11 | 21,76 | 0,357 | 253,6 | 8,65 | | |
| *Grewia fallax* | GRFA | 2,17 | 21,89 | 0,439 | 234,8 | 8,82 | 0,215 | 0,469 |
| *Kigelia africana* | KIAF | 1,56 | 26,22 | 0,303 | 323,3 | 7,95 | 0,237 | 0,330 |
| *Ormocarpum trichocarpum* | ORTR | 3,54 | 12,08 | 0,322 | 238,9 | 12,36 | 0,175 | 0,590 |
| *Sclerocarya birrea* | SCBI | 2,04 | 22,57 | 0,393 | 281,0 | 7,78 | 0,228 | 0,440 |
| *Senegalia polyacantha* | SEPO | 3,65 | 13,22 | 0,456 | 129,2 | 14,15 | 0,124 | 0,587 |
| *Senegalia senegal* | SESE | 4,71 | 9,91 | 0,488 | 176,3 | 12,93 | 0,170 | 0,577 |
| *Terminalia mollis* | TEMO | 1,03 | 46,90 | 0,461 | 338,4 | 5,39 | 0,190 | 0,520 |
| *Vachellia drepanolobium* | VADR | 3,44 | 13,63 | 0,483 | 154,5 | 12,71 | 0,124 | 0,624 |
| *Vachellia gerrardii* | VAGE | 2,15 | 22,15 | 0,463 | 242,2 | 9,00 | 0,161 | 0,553 |
| *Vachellia hockii* | VAHO | 2,98 | 16,24 | 0,490 | 121,3 | 14,62 | 0,202 | 0,556 |
| *Vachellia nilotica* | VANI | 1,95 | 25,06 | 0,547 | 184,3 | 9,98 | 0,165 | 0,606 |
| *Vachellia robusta* | VARO | 2,23 | 21,83 | 0,503 | 237,8 | 7,88 | 0,158 | 0,577 |
| *Vachellia seyal* | VASE | 3,34 | 13,03 | 0,352 | 168,3 | 16,26 | 0,139 | 0,543 |
| *Vachellia sieberiana* | VASI | 2,44 | 18,77 | 0,503 | 192,2 | 10,30 | 0,150 | 0,445 |
| *Vachellia tortilis* | VATO | 3,33 | 14,34 | 0,472 | 131,3 | 14,85 | 0,137 | 0,592 |
| *Ziziphus mucronata* | ZIMU | 1,25 | 38,92 | 0,369 | 340,1 | 7,59 | | |

Note: Bark investment (Bark inv) gives the slope of the relationship between the diameter of the stem (cm) and bark thickness (cm). Height investment (height inv) gives the slope of the relationship between the diameter of the stem (cm) and vertical height (m). *Average weight for 1000 seeds (g) (Liu et al. 2019).

Table S2 (continued)

| **Species** | **Code** | **Seed mass (g*)** | **Max height (m)** | **Max RD (ind 100 $^{m-2}$)** | **TDMC** | **Mech defense** |
|---|---|---|---|---|---|---|
| *Albizia harveyi* | ALHA | 87,00 | 15 | 1,08 | 0,544 | low |
| *Balanites aegyptiaca* | BAAE | 1377,78 | 12 | 0,14 | 0,507 | low |
| *Boscia angustifolia* | BOAN | 52,33 | 10 | | | low |
| *Commiphora africana* | COAF | 109,16 | 10 | | 0,345 | low |
| *Combretum molle* | COMO | 89,60 | 7 | 9,95 | 0,511 | low |
| *Cordia ovalis* | COOV | 97,33 | 8 | | 0,539 | low |
| *Commiphora trothea* | COTR | 124,97 | 6 | 5,42 | 0,385 | low |
| *Dichrostachys cinerea* | DICI | 19,56 | 8 | 21,63 | 0,400 | low |
| *Euclea natalensis* | EUNA | 147,30 | 12 | | | low |
| *Grewia bicolor* | GRBI | 112,30 | 6 | | | low |
| *Grewia fallax* | GRFA | | 9 | 1,68 | 0,445 | low |
| *Kigelia africana* | KIAF | 144,11 | 18 | | 0,387 | low |
| *Ormocarpum trichocarpum* | ORTR | 12,45 | 5 | 52,92 | 0,365 | low |
| *Sclerocarya birrea* | SCBI | 2000,00 | 18 | | 0,460 | low |
| *Senegalia polyacantha* | SEPO | 90,00 | 21 | | 0,441 | high |
| *Senegalia senegal* | SESE | 72,00 | 10 | 9,80 | | high |
| *Terminalia mollis* | TEMO | 756,00 | 13 | | 0,39 | low |
| *Vachellia drepanolobium* | VADR | 37,41 | 5 | 17,83 | | high |
| *Vachellia gerrardii* | VAGE | 78,97 | 15 | 6,82 | 0,460 | high |
| *Vachellia hockii* | VAHO | 32,10 | 6 | 5,88 | 0,454 | high |
| *Vachellia nilotica* | VANI | 148,83 | 14 | 2,32 | 0,550 | high |
| *Vachellia robusta* | VARO | 91,00 | 25 | 2,39 | 0,466 | low |
| *Vachellia seyal* | VASE | 46,66 | 17 | 1,29 | 0,432 | high |
| *Vachellia sieberiana* | VASI | 333,00 | 18 | 7,17 | 0,517 | high |
| *Vachellia tortilis* | VATO | 58,06 | 21 | 2,38 | 0,513 | high |
| *Ziziphus mucronata* | ZIMU | 864,11 | 15 | | 0,364 | low |
